# The MIME-seq technique allows to monitor the interaction of small non-coding RNAs with Argonaute proteins and their transfer to other cells

**DOI:** 10.64898/2025.12.18.695118

**Authors:** Jérôme Perrard, Claudiane Guay, Nadège Zanou, Romano Regazzi

## Abstract

The MIME-seq2.0 technique utilizes a chimeric HENMT1 methyltransferase engineered to bind to Argonaute proteins (hereafter referred to MIME enzyme) to protect the terminal sugar of small non-coding RNAs (sncRNAs) from oxidation, a reaction that inhibits unmodified RNA ligation and cloning. In this study, we confirm that the MIME-seq technique efficiently distinguishes oxidized and non-oxidized sncRNAs. Indeed, expression of the MIME enzyme in the insulin-secreting cell line MIN6B1 protected miRNAs from oxidation and permitted their detection by RNA sequencing and quantitative PCR. Comparison of the sncRNA profile between wild type and MIME-expressing cells demonstrated protection from oxidation not only of miRNAs, but also of some Y-RNA and tRNA-derived fragments. Immunoprecipitation with Ago2 antibodies confirmed that the protected Y-RNA and tRNA fragments bind to Argonaute proteins. We also used this system to track miRNA transfer between cells via extracellular vesicles (EVs). miRNAs released inside EVs of MIME-expressing Jurkat T cells or C2C12 myotubes and delivered to MIN6B1 cells were protected from oxidation, enabling sensitive and specific detection of the transferred miRNAs in the receiving cells. Overall, the MIME-seq technique provides a powerful tool for analyzing small RNA binding to Argonaute proteins in living cells and offers new possibilities for tracking RNA transfer across different cell types.

## INTRODUCTION

The human genome consists of approximately 3.2 billion base pairs, but less than 2% encode proteins. The majority of transcribed genomic sequences give rise to a diverse array of non-coding RNAs, which can be classified based on length, physicochemical properties, genomic origin, or binding partners [1]. These non-coding RNAs include microRNAs (miRNAs), Piwi-interacting RNAs (piRNAs), long non-coding RNAs, and circular RNAs. Additionally, mammalian cells contain thousands of small non-coding RNAs (sncRNAs) derived from larger transcripts, such as tRNA-derived fragments (tRFs), rRNA-derived fragments, fragments derived from Y RNAs and small intronic RNAs (sinRNAs) [1].

Among sncRNAs, miRNAs are the most well-studied, playing a crucial role in gene regulation. The biogenesis and function of these 21-23 nucleotide molecules have been extensively investigated [2]. Mature miRNAs are generated from precursor miRNAs by cleavage by the endonuclease Dicer and subsequently bind to the Argonaute 2 (Ago2) protein, forming part of the RNA-Induced Silencing Complex (RISC). This association allows the RISC complex to bind complementary sequences in the 3’ untranslated region (UTR) of target mRNAs, leading to the repression of their translation into proteins. Other sncRNAs, including tRFs, piRNAs, and sinRNAs, have been proposed to interact with Ago2 and to function as translational repressors [3, 4]. However, these interactions have primarily been suggested through co-immunoprecipitation studies with Ago2, and the direct involvement of these small RNAs in RISC-mediated regulation in intact cells remains largely unconfirmed.

In addition to their intracellular roles, several sncRNAs are secreted inside extracellular vesicles (EVs) or associated to RNA-biding proteins and can be delivered to nearby or distant cells [5]. This extracellular signaling has been proposed to play an important role in intercellular and interorgan communication [6]. Despite growing interest in this novel form of communication, sensitive and reliable methods for monitoring the intercellular exchange of miRNAs and other sncRNAs remain limited.

The MIME-seq2.0 method, developed by Mandlbauer et al. [7], was designed to detect cell-specific miRNA pools in complex tissues without the need for cell isolation. This technique leverages a chimeric HENMT1 enzyme engineered to bind Argonaute proteins instead of Piwi proteins that methylates miRNAs at their 3’ terminal 2’-OH [7]. Unmethylated miRNAs are oxidized by sodium periodate (NaIO_4_), preventing their ligation to sequencing adapters, while methylated miRNAs are resistant to oxidation and can be efficiently detected by RNA sequencing. In mammalian cells, miRNA methylation occurs only when the methyltransferase is engineered to bind to Argonaute (Ago) proteins [7]. Since RNA methylation by chimeric HENMT1 depends on Ago binding, we reasoned that this method could identify, in an unbiased manner, the pool of sncRNAs potentially associated with the RISC complex.

In this study, we applied the MIME-seq2.0 technique to gain a comprehensive picture of the sncRNAs associated with Argonaute proteins. In addition to miRNAs, we identified a group of tRFs and Y RNA-derived fragments that are methylated by chimeric HENMT1 and co-immunoprecipitated with Ago2. Furthermore, we demonstrate that MIME-seq2.0 can efficiently track the transfer of miRNAs between cells, offering new insights into this emerging form of cellular communication.

## MATERIALS AND METHODS

### Cell Culture

The MIN6B1 cell line [8] was cultured in DMEM-GlutaMAX medium supplemented with 15% decomplemented FBS, 70 µM β-Mercaptoethanol, 50 mg/mL streptomycin, and 50 IU/mL penicillin. The Jurkat clone 20 cell line was obtained from O. Acuto (Rebeaud et al., 2008). Jurkat T cells were cultured in RPMI 1640 medium supplemented with 10% decomplemented FBS, 50 mg/mL streptomycin, and 50 IU/mL penicillin. HEK293T cells and C2C12 cells were obtained from the American Type Culture Collection (ATCC-ACS-4500 and ATCC-CRL-1772, respectively). Both cell lines were cultured in DMEM-GlutaMAX medium containing 10% decomplemented FBS, 50 mg/mL streptomycin, and 50 IU/mL penicillin. All cells were maintained at 37°C in a humidified atmosphere with 5% CO2 and were tested negative for mycoplasma contamination.

### Cell transfection and transduction

The plasmid for lentivirus production and MIME enzyme expression was kindly provided by Dr. Luisa Cochella, Johns Hopkins University, Baltimore. FLAG-Ago2 was a gift from Edward Chan (Addgene plasmid # 21538). Nucleic acids were transfected using Lipofectamine 2000 according to the manufacturer’s protocol. Cells were incubated with the transfection complex in antibiotic-free medium for 4 hours, followed by incubation in regular media for 24 to 72 hours. Lentivirus was produced by transfecting 1.2 µg of the MIME enzyme-expressing plasmid, 0.4 µg of pCMVR8.74, and 0.4 µg of pVSV-G into HEK293T cells for 48 hours. The culture medium was filtered using 0.45 µm filters and used to infect C2C12 cells in the presence of Polybrene (8 µg/µL), or Jurkat cells via spinoculation (800 g, 30 minutes at RT, in the presence of Polybrene, 8 µg/µL). Infected C2C12 or Jurkat cells were selected with Blasticidin (5 µg/mL for C2C12 and 10 µg/mL for Jurkat).

### Protein extraction and immunoprecipitation

MIN6B1 cells transfected using control or Flag-Ago2 plasmids were grown in 10 cm plates, detached using trypsin, washed, pelleted, and stored as dried pellets at -80°C until needed. The pellets were thawed on ice and homogenized in 500 µL of lysis buffer (1% Triton X-100, 150 mM NaCl, 10 mM Tris pH 7.5, 1 mM EDTA), supplemented with a protease inhibitor cocktail, 0.16 U/µL RNase inhibitor, and 0.5 mM DTT by pipetting. After incubating for 15 minutes, lysates were centrifuged at 16,000xg for 10 minutes at 4°C. Protein concentrations were determined by Bradford assay. Equal amounts of protein extracts were incubated with 20 µL equilibrated anti-Flag magnetic beads (Sigma M8823) for 2 hours at 4°C under agitation. Beads were isolated using a magnet and washed eight times (five washes with lysis buffer and three washes with PBS). One fifth of the beads was incubated in 1X Laemmli buffer to measure protein content while the rest was used for RNA extraction.

### Western blotting

Protein samples were heated at 95°C for 5 minutes, then loaded onto 4-20% gradient SDS-PAGE gels (Bio-Rad) and transferred to a nitrocellulose membrane using a wet transfer system at 100 V. The membrane was blocked using 5% BSA solution in PBST and incubated with primary antibody against Myc tag (c-Myc 9E10 clone), Flag tag (Sigma-Aldrich F3165) or Actin (Sigma-Aldrich), followed by an additional incubation step with HRP-coupled anti-Mouse secondary antibody (Biorad).

### Production and isolation of extracellular vesicles

EV-free media was prepared by centrifuging 30% FBS-containing media at 100,000xg for 2 hours at 4°C. The supernatant was diluted with fresh media to match the normal FBS concentration and filtered-sterilized. Jurkat T cells were seeded at a density of 0.1–0.3 million cells/mL and cultured in EV-free media for 3 days. C2C12 cells were induced to differentiate by replacing the culture media with differentiation media (2% Horse Serum supplemented with non-essential amino acids and 50 mg/mL streptomycin, 50 IU/mL penicillin) once they were fully confluent. Differentiation media was changed every 2–3 days. After 5 days of differentiation, the media was replaced with EV-free differentiation media and was harvested 2 and 5 days later.

For all experiments, the collected media was immediately centrifuged at 300xg for 5 minutes, then at 2000xg for 10 minutes, and 10,000xg for 30 minutes at 4°C. The supernatant was frozen at - 20°C before further processing. The supernatant was then filtered through a 0.45 µm filter and ultracentrifuged at 100,000xg for 2 hours at 4°C. The EV pellets were washed with PBS and centrifuged again under the same conditions. A final wash was performed using the same parameters in a Sorvall WX 80 centrifuge with the TST604 rotor. An aliquot of the supernatant was taken for analysis. The EV pellets were resuspended in 100 µL of supernatant and stored at - 80°C for further use. Particle concentration was assessed using a ZetaSizer.

### Treatment with extracellular vesicles

MIN6 cells were seeded at 105,000 cells/cm^2^ in 24-well plates. The following day, the media was replaced with EV-free media, and 100 µL of PBS, SN, or EV preparation was added to the wells for 24 hours. The EV concentration ranged from 0.5 to 5 × 10^10^ particles of approximately 150 nm per milliliter, depending on the experimental replicate. After 24 hours, cells were harvested following two PBS washes, using Qiazol reagent, and stored at -20°C until further processing.

### RNA extraction

Total RNA, including small RNAs, was extracted using the miRNeasy Micro Kit (Qiagen) according to the manufacturer’s instructions. Total RNA was eluted with water, and RNA concentration was determined using a Nanodrop 1000.

### MIME-seq technique

The oxidation step for MIME-seq was performed as previously described [7]. Briefly, equal amounts of RNA for each condition were diluted with water and borate buffer (60 mM final concentration). A 2 µL aliquot of NaIO_4_ (2.5 mM final) or water was added to 18 µL of the RNA mixture and incubated at room temperature for 30 minutes. The reaction was stopped by adding 5 volumes of water, NaOAc (300 mM final, pH 5.2), 20 µg of Glycogen, and 300 µL of ice-cold ethanol. The samples were incubated at -80°C for at least one hour, then centrifuged at 20,000xg for 30 minutes at 4°C. The RNA pellets were washed with 80% ethanol, centrifuged briefly, dried, and resuspended in pure water.

### Small RNA-seq

Libraries were prepared using the QIAseq miRNA Library Kit (Qiagen) according to the manufacturer’s instructions and analyzed using the Agilent BioAnalyzer 2100. The samples were then sequenced on an Illumina NextSeq 500 instrument. Raw data analysis was performed using the Qiagen RNA-seq Analysis Portal to generate miRNA and piRNA lists, and additional analysis was carried out by the Bioinformatics Competence Center to generate an unbiased list of small RNA sequences [9].

### RT-qPCR

After oxidation, equal volumes of purified RNA (typically containing 200 ng) were used for reverse transcription using the miRCURY LNA RT Kit (Qiagen). The kit includes a critical ligation step to distinguish oxidized RNA from non-oxidized RNA. After a 1-hour incubation at 42°C and 5-minute incubation at 95°C, cDNA was diluted 6-12 times, and 4 µL of the dilution was used for qPCR using the miRCURY LNA SYBR Green PCR Kit. Probes for Let-7b-5p, miR-1-3p, miR-7-5p, miR-16-5p, miR-133b, miR-142-3p, miR-142-5p, mmu-miR-155-5p, hsa-miR-155-5p, U6, Y1, Y3, and Val-CAC were used (Supplementary Table 1). Samples with no amplification (>50 Ct) or aberrant melting curves were excluded from analysis. qPCR results were analyzed using the formula “Remaining miRNA = 100 / (2^ΔCt)” (oxidized vs. water condition) to calculate the remaining non-oxidized RNA after the oxidation step. Negative controls with water instead of samples showed no amplification.

## RESULTS

The MIME-seq2.0 technique developed by Mandlbauer et al. [7] employs the HENMT1ΔC-T6B methyltransferase, hereafter referred to as the MIME enzyme, which is engineered to bind Argonaute proteins instead of Piwi proteins (Fig.1A). By methylating the terminal sugar, this enzyme protects the target RNA from oxidation by sodium periodate (Oxidation Step, Fig.1B). In unmethylated RNAs the oxidation impedes ligation and cloning but does not affect hybridization. To distinguish between oxidized and non-oxidized species, a ligation step is required, such as the one used in RNA sequencing techniques. We hypothesized that the ligation step during reverse transcription of sncRNAs using the miRCURY LNA RT Kit would also allow differentiation of oxidized from non-oxidized species by qPCR. To test this, we transiently transfected plasmids expressing or lacking the MIME enzyme into MIN6B1 cells, a mouse insulinoma cell line. Western blot analysis confirmed robust expression of the MIME enzyme in MIN6B1 cells, as detected by anti-Myc antibody (Fig. 1C). Following NaIO_4_ oxidation of total RNA from both transfected and non-transfected cells, qPCR analysis showed almost complete loss (>10 ΔCt) in the oxidized condition of let-7b-5p and miR-7-5p, two among the most abundant miRNAs in MIN6B1 cells. Indeed, after oxidation the recovery rate of these miRNAs was <0.1%. In contrast, in cells expressing MIME, let-7b and miR-7 miRNAs were easily detected even after oxidation, yielding a recovery rate of ∼25% (2 ΔCt difference). U6, a non-Argonaute-interacting sncRNA, showed a recovery rate of ∼1% regardless of MIME enzyme presence. These results demonstrate that this approach can effectively assess the methylation status of small RNAs via qPCR.

**Fig 1.**
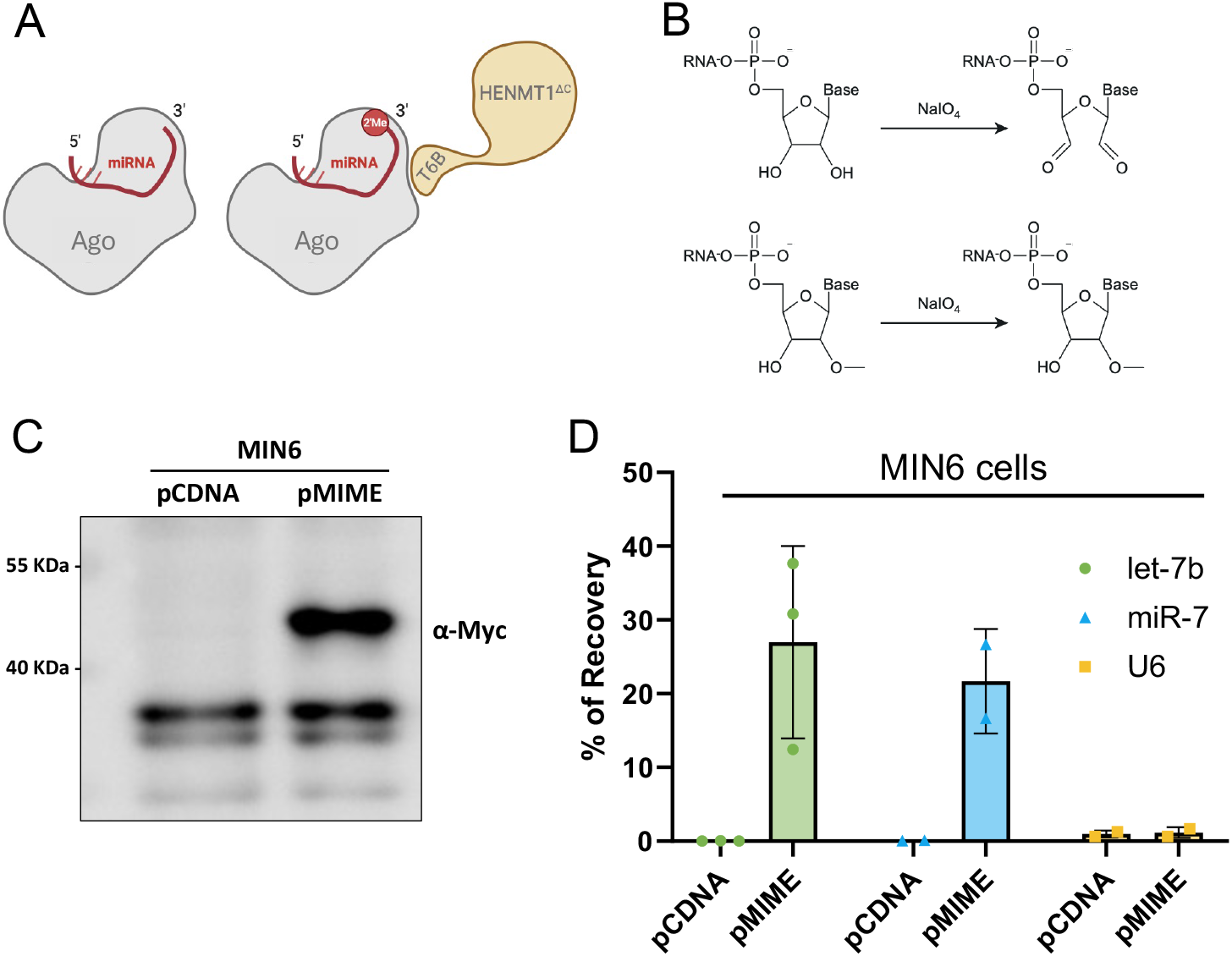
Application of the MIME-seq 2.0 technique to insulin-secreting cells. **A)** Design of the chimeric HENMT1 enzyme binding to Argonaute proteins. **B)** Effect of NaIO_4_ treatment on unmethylated (upper row) and methylated (lower row) terminal nucleotide. **C)** MIN6B1 cells were transfected with a control plasmid (pCDNA) or with a plasmid encoding a myc-tagged MIME enzyme. The expression of MIME was assessed by western blotting using an antibody against myc. **D)** RNA extracted from MIN6B1 cells transfected with pCDNA or with the plasmid encoding the MIME enzyme was treated with NaIO4 as indicated in the Methods section. The expression of the indicated sncRNAs was measured by qRT-PCR. The data are expressed as mean ± SEM (n=3).

Because U6 is not methylated by HENMT1ΔC-T6B and the enzyme cannot methylate miRNAs in the absence of its T6B Argonaute-binding region [7], we hypothesized that this system could also be used to evaluate sncRNA binding to Argonaute proteins. To test this, we performed MIME-seq to analyze small RNA methylation status in the presence or absence of the MIME enzyme (GEO accession number GSE313744, reviewers token upwtwyoybnutpuh). RNA sequencing was initially analyzed using the Qiagen RNA-seq Analysis Portal to assess miRNA and piRNA expression (Supplementary Table 2), followed by unbiased analysis of other sncRNAs (Supplementary Table 3).

Principal component analysis (Fig. 2A) showed that in unoxidized conditions, the RNA profiles were closely clustered regardless of MIME enzyme expression. However, the oxidation step in the presence of MIME caused a slight shift in the principal components, while the oxidation step without MIME significantly altered the profile. As expected, miRNA recovery was minimal under basal conditions, indicating they are not methylated by endogenous enzymes in cellulo (Fig. 2B, Supplementary Table 2). In contrast, the MIME enzyme provided robust protection to most miRNAs. Many piRNAs are known to be methylated at the 3’ terminal 2’-OH end by endogenous HENMT1 [10]. Thus, as expected, many piRNAs (e.g., mmu-piR-67265, mmu-piR-68892) were protected from oxidation even without MIME (Supplementary Table 2). However, consistent with the observation that not all piRNAs are terminally methylated, some piRNAs (e.g., mmu-piR-69027, mmu-piR-67787) were not protected from oxidation. Notably, MIME expression did not change the protection levels of piRNAs, suggesting that they do not associate to Ago proteins and are not accessible to chimeric HENMT1 lacking the Piwi-interacting domain (Fig. 2B).

**Fig 2.**
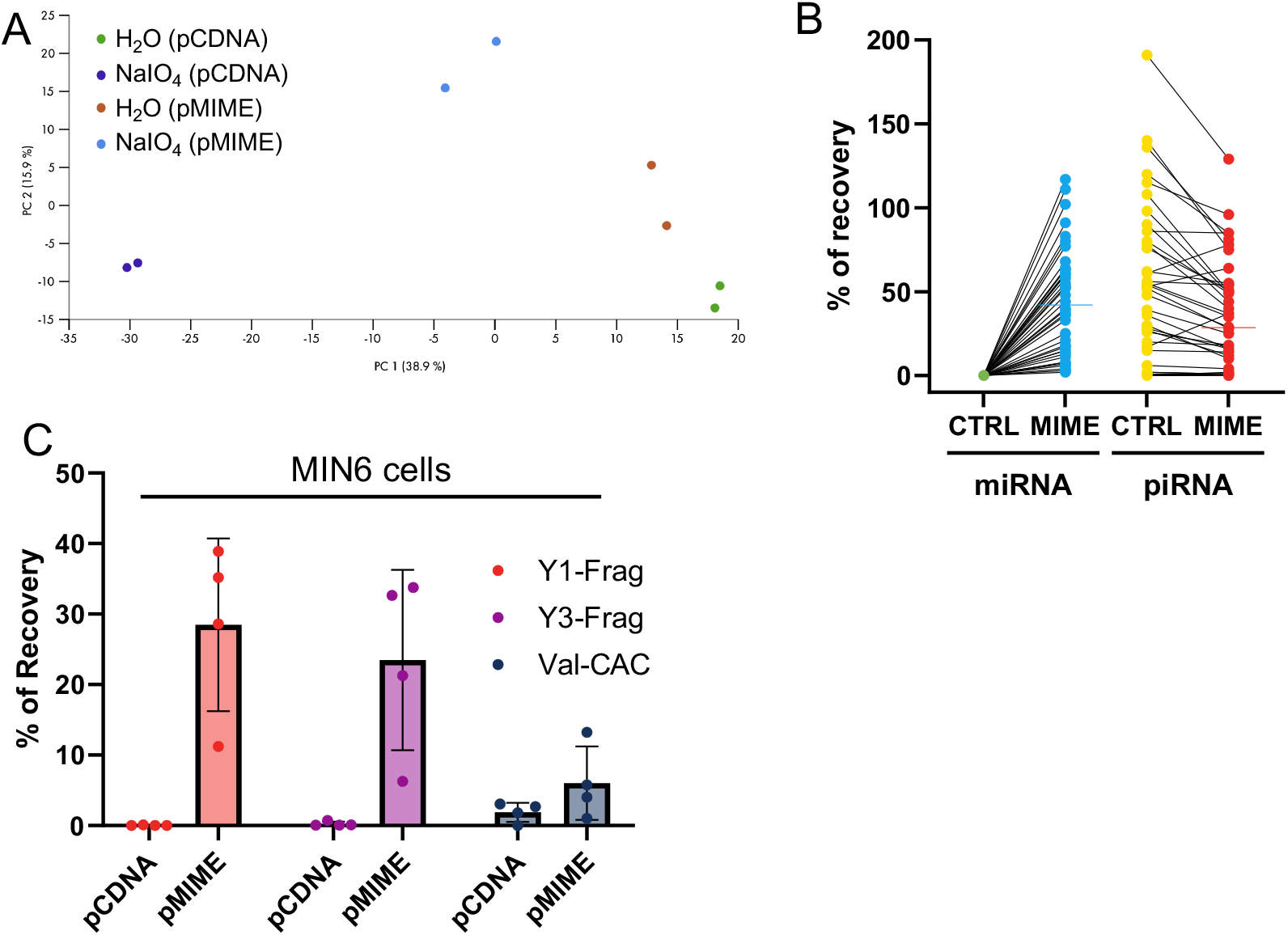
Comprehensive analysis of the sncRNAs detectable using the MIME-seq approach. RNA samples obtained from MIN6 cells expressing or not the MIME enzyme were incubated in the presence or absence of NaIO4. Samples from two independent experiments were then analyzed by small RNA sequencing. **A)** Principal component characteristics of the sequencing data. **B)** Recovery rates of miRNAs and piRNAs after NaIO_4_ treatment in samples obtained from MIN6 cells expressing or not the MIME enzyme. The data are expressed as percent of the level measured in cells not exposed to NaIO_4._ **C)** Confirmation by qRT-PCR of the recovery rate of the indicated sncRNAs.

Beside miRNAs, unbiased analysis revealed partial protection by the MIME enzyme of other small non-coding RNAs, such as Y1 and Y3 fragments from the Y-RNA family, as well as tRFs like tRF^Val-CAC^ and tRF^Asp-GTC^ (Supplementary Table 3). Protection of these fragments was confirmed by qPCR, with Y1 and Y3 fragments showing recoveries similar to that of miRNAs, while the recovery of tRF^Val-CAC^ was less efficient (from 1.9% to 6.0% with MIME, Fig. 2C).

To confirm the binding of these RNA fragments to Argonaute proteins, we performed RNA immunoprecipitation (RNA-IP) using Flag-Ago2. The expression and enrichment of Ago2 following anti-Flag immunoprecipitation were confirmed by Western blot (Fig. 3A). miRNA binding to Flag-Ago2 was validated by recovery percentages of let-7b-5p and miR-7-5p (5.7% and 4.1%, respectively) in the Flag-Ago2 IP, with negligible recovery in control IP (0.06% and 0.03%). Interestingly, Y1 fragments were highly enriched in the Flag-Ago2 IP (1.9%) compared to the control (0.004%), indicating significant binding to Ago2 (Fig. 3B). Y3 and tRF^Val-CAC^ fragments were also co-immunoprecipitated by Ago2, though to a lesser extent than miRNAs (Fig. 3C). As expected, U6 was not enriched in the Ago2 IP. These results suggest that MIME-seq can be used to evaluate sncRNA binding to Argonaute proteins.

**Fig 3.**
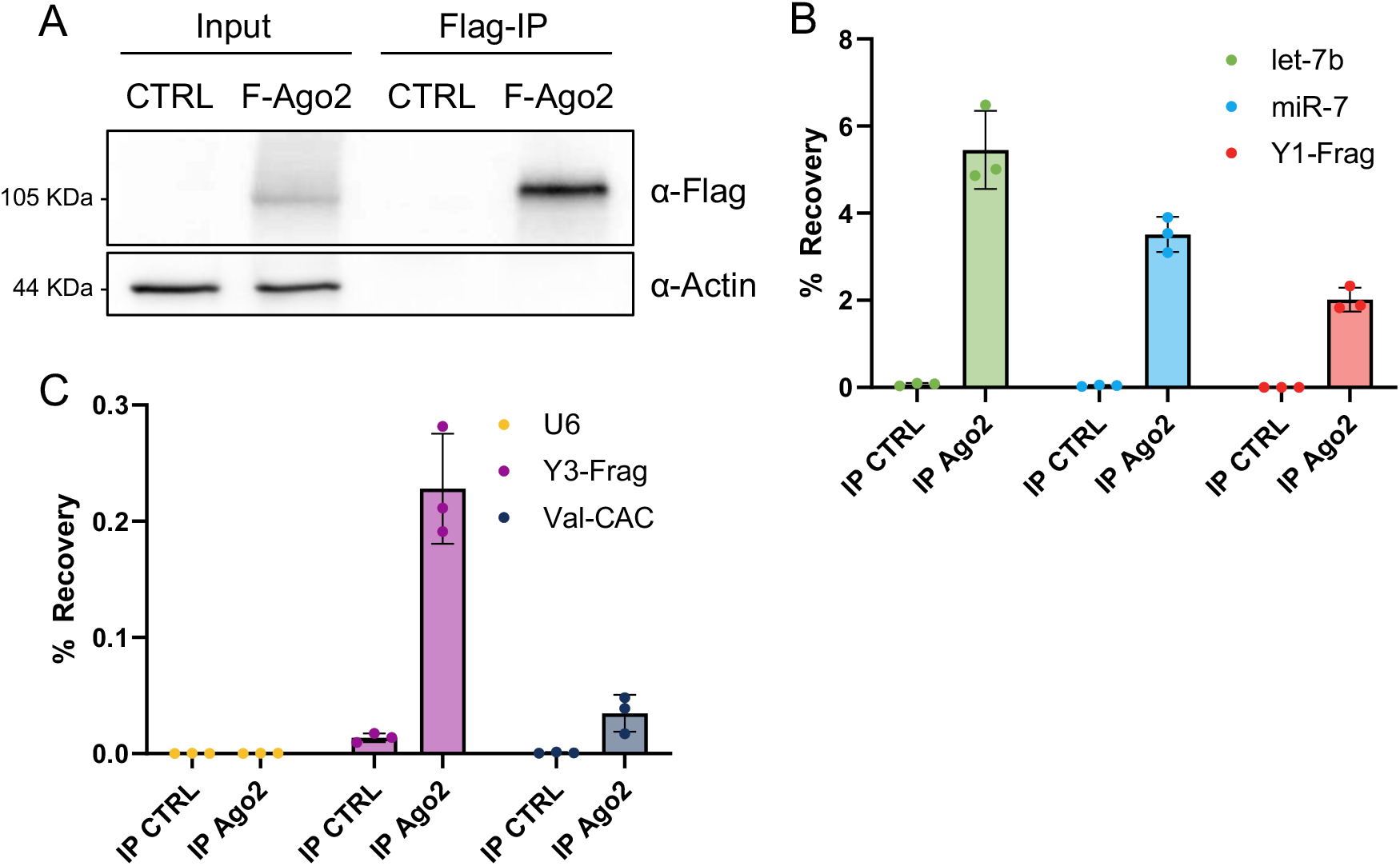
Confirmation of the binding of different sncRNAs to Ago 2. **A)** Samples of control (CTRL) or Flag-Ago2 transfected MIN6 cells were incubated in the presence of anti-FLAG antibody-coupled beads. The level of Ago2 in input and after immunoprecipitation was assessed by Western blotting. The level of α-Actin was measured in parallel as loading control. **B and C)** Recovery of the indicated sncRNAs after immunoprecipitation with anti-Flag beads of control (IP CTRL) or Flag-Ago2 (IP Ago2) transfected samples was assessed by qRT-PCR.

Given the ability of the MIME enzyme to label small RNAs bound to Argonaute proteins, we hypothesized that this technique could be used to track miRNA transfer between cells with high specificity and sensitivity (Fig.4A). To test this, we used a model of miRNA transfer via extracellular vesicles (EVs) from human Jurkat CD4^+^ T cells to mouse MIN6 cells [11]. Jurkat T cells were transduced to express the MIME enzyme. As expected, miRNAs from MIME-expressing Jurkat T cells were protected from oxidation, while miRNAs from wild-type (WT) Jurkat T cells were not (Fig. 4B). For example, recovery rates for miR-7 and miR-142-3p were 41% and 32% in MIME-expressing Jurkat T cells, but only 0.57% and 0.09% in WT cells, respectively. miR-142-5p was also protected in the presence of the MIME enzyme, although slightly less efficiently (5.7% vs. 0.07% in WT). In contrast, U6 recovery was unaffected by the MIME enzyme.

**Fig 4.**
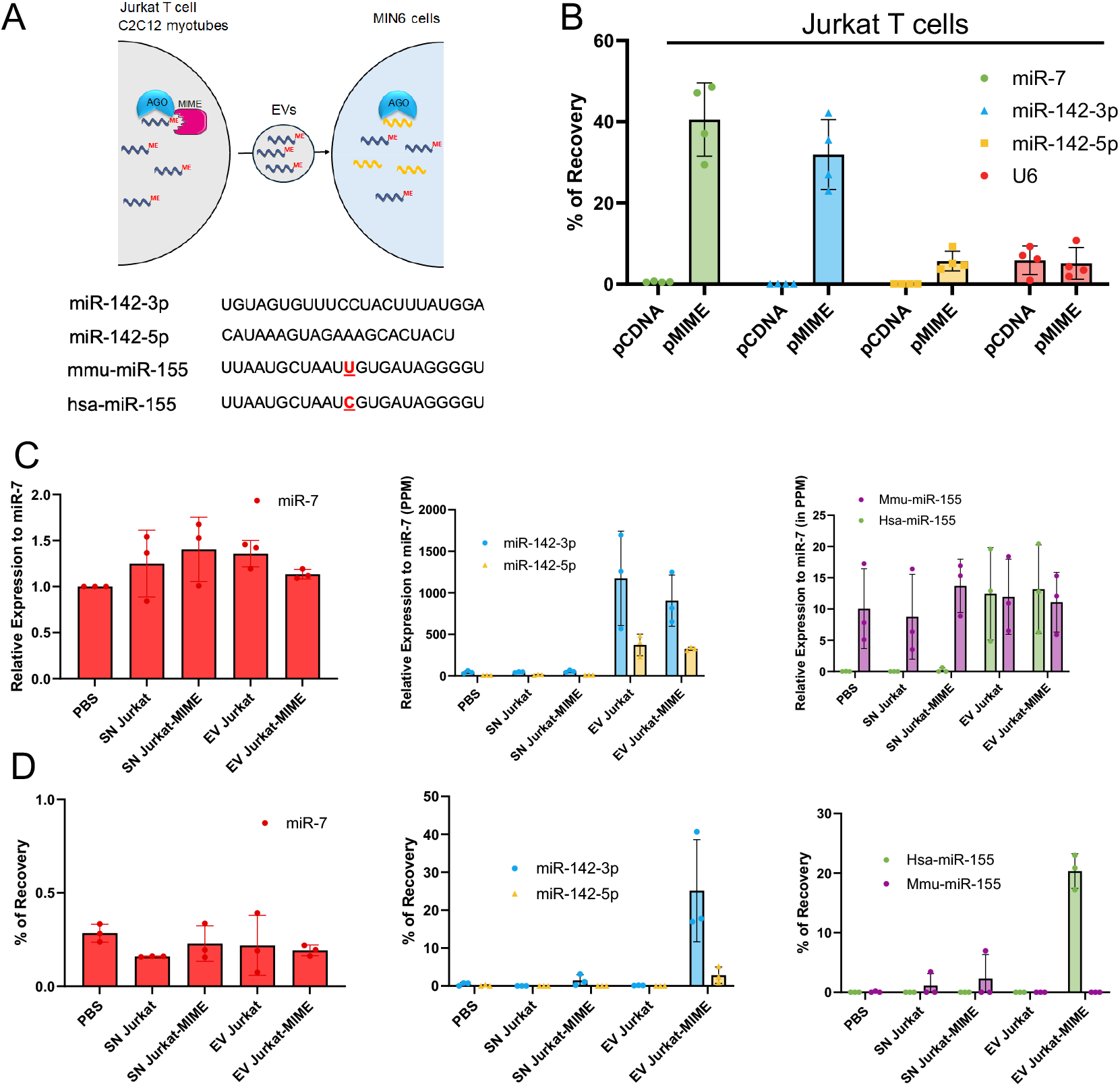
Use of the MIME-seq technique to monitor the transfer of miRNAs from human Jurkat T cells to mouse insulin-secreting MIN6B1 cells. **A)** Sequences of the measured miRNAs. **B)** Recovery of the indicated sncRNAs in Jurkat T cells expressing or not the MIME enzyme after the oxidation step. The data show the results of four independent experiments. **C)** Measurement of the level of the indicated miRNAs in MIN6B1 cells incubated in the presence or absence of EVs produced by wild type Jurkat T cells or Jurkat T cells expressing the MIME enzyme. **D)** Recovery rate after NaIO_4_ treatment of the indicated miRNAs in MIN6B1 cells incubated in the presence or absence of EVs produced by wild type Jurkat T cells or Jurkat T cells expressing the MIME enzyme.

Next, we purified by differential ultracentrifugation EVs released from the human Jurkat T cell line expressing or not the MIME enzyme. Without the oxidation step, miR-142-3p, miR-142-5p, and human miR-155-5p (hsa-miR-155-5p) were upregulated in murine MIN6B1 insulinoma cells upon incubation with EVs from both transduced and WT Jurkat cells (Fig. 4C), confirming previous findings [11]. As expected, the level of mouse miR-155-5p (mmu-miR-155-5p), which is endogenously produced by MIN6B1 cells and differ by one nucleotide from hsa-miR-155-5p (Fig.4A), was unaffected by EVs treatment. After the oxidation step, only the cells incubated with EVs from MIME-expressing Jurkat cells showed significant recovery of miR-142-3p, miR-142-5p and hsa-miR-155-5p, while endogenously produced mmu-miR-155-5p was not protected (Fig. 4D). These results demonstrate that the rise in the levels of miR-142-3p, miR-142-5p and miR-155-5p observed after incubation of EVs from Jurkat CD4^+^ T cells results from the transfer of these sncRNAs and is not due to increased expression in the receiving cells.

To further validate this technique, we tested it in a different cell model: mouse C2C12 myotubes. After inducing C2C12 differentiation and collecting EVs from myotubes, we incubated MIN6B1 insulinoma cells with these EVs. MiR-16-5p and miR-7-5p expression was unaffected by the EV treatment, most likely because the amount of these miRNAs transferred via myotube EVs is too small compared to the endogenous level in MIN6B1 cells. In contrast, miR-1-3p and miR-133b, two miRNAs highly expressed in myotubes [12], were upregulated in MIN6B1 cells incubated with EVs from both WT C2C12 cells and MIME-expressing C2C12 cells (Fig. 5A). After oxidation, miR-1 and miR-133b were recovered only in EVs from transduced C2C12 cells (Fig. 5B). This confirms that MIME-seq can be used to track miRNA transfer between different cell lines and tissues.

**Fig 5.**
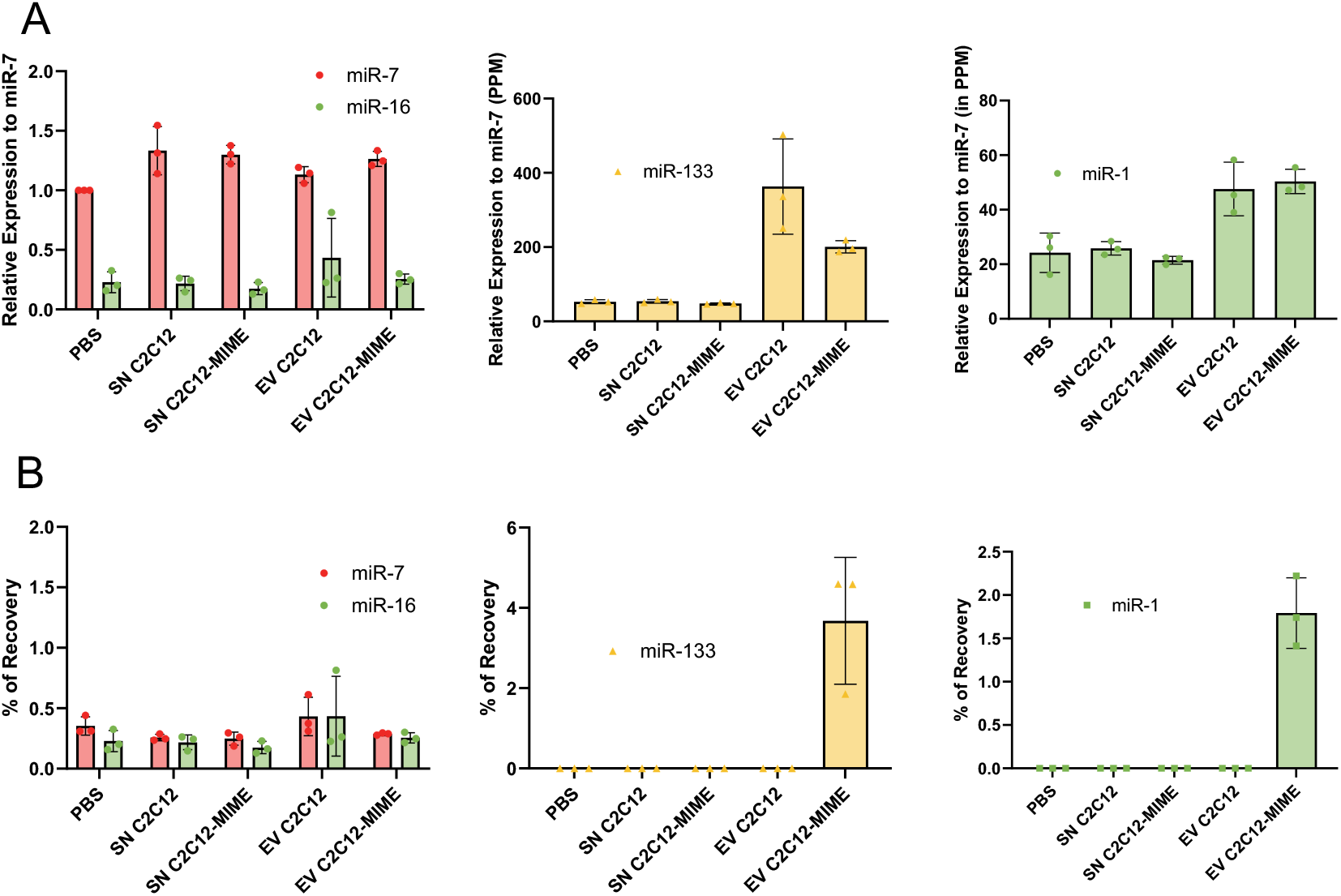
Use of the MIME-seq technique to monitor the transfer of miRNAs from C2C12 myotubes to insulin-secreting MIN6B1 cells. **A)** Measurement of the level of the indicated miRNAs in MIN6B1 cells incubated in the presence or absence of EVs produced by wild type C2C12 myotubes or C2C12 myotubes expressing the MIME enzyme. **B)** Recovery rate after NaIO_4_ treatment of the indicated miRNAs in MIN6B1 cells incubated in the presence or absence of EVs produced by wild type C2C12 myotubes or C2C12 myotubes expressing the MIME enzyme.

## DISCUSSION

The results presented here demonstrate the utility of MIME-seq 2.0, a technique utilizing the HENMT1ΔC-T6B methyltransferase (MIME enzyme), in distinguishing oxidized from non-oxidized sncRNAs. This method offers a novel approach for assessing the methylation status of miRNAs and other sncRNAs and for evaluating their interactions with Ago proteins. This information would be instrumental in assessing the possible mode of action of sncRNAs of unknown function. Furthermore, we show that MIME-seq 2.0 can be applied to track miRNA transfer between cells, including through EVs, providing a tool for high-specificity, high-sensitivity studies of RNA dynamics.

In agreement with the observations of Mandlbauer et al. [7], we found that the MIME enzyme efficiently protects small RNAs, particularly miRNAs, from oxidation. The ability of MIME to maintain the integrity of these RNAs after oxidation, with ∼25% recovery rates for let-7b and miR-7, is striking compared to the ∼0.1% recovery in the absence of MIME. Importantly, this protection is specific to methylated species, since recovery of U6 RNA which is not methylated by the MIME enzyme is negligible. The strong differential recovery rates observed in RNA-seq and qPCR assays following oxidation demonstrate that MIME-seq 2.0 can effectively discriminate between oxidized and non-oxidized RNA species.

The potential for MIME-seq to analyze RNA binding to Argonaute proteins was also explored, particularly in light of the enzyme’s selective methylation of miRNAs and other sncRNAs that interact with Argonaute proteins. The principal component analysis of RNA sequencing data demonstrated that the expression of MIME caused only subtle shifts in the RNA profiles under oxidized conditions, consistent with the fact that the enzyme selectively protects certain species from oxidation. The absence of MIME, however, resulted in significant disruption of the RNA profile, indicating that oxidation without protection substantially alters the RNA landscape.

Interestingly, while the majority of the miRNAs were protected by MIME in oxidized conditions, many piRNAs were already methylated and thus naturally protected from oxidation, regardless of MIME expression. Notably, MIME expression did not improve the recovery of the pool of piRNAs affected by the oxidation step, reinforcing the notion that only small RNAs associated to Ago proteins can be methylated by the MIME enzyme. In contrast, we observed protection of other classes of sncRNAs, including Y-RNA fragments (Y1 and Y3) and a group of tRFs, indicating that the MIME enzyme can protect a wider range of small RNAs, although with varying efficiency. These findings were further supported by our RNA immunoprecipitation data showing Ago2 enrichment for the protected sncRNAs. Our findings are consistent with previous reports showing that Y-RNA fragments and some tRFs can associate to Ago proteins [3]. This highlights the broad applicability of MIME-seq to assess the interaction of different sncRNA families to Argonaute proteins.

A major advantage of MIME-seq 2.0 resides in its potential for tracking the transfer of small RNAs between cells, particularly through EVs. Our experiments demonstrated that miRNAs transferred via EVs from Jurkat T cells expressing MIME were efficiently protected from oxidation, whereas miRNAs from wild-type Jurkat T cells and those endogenously produced by the receiving cells were not. This observation confirms that the MIME enzyme labels transferred miRNAs with methyl groups, allowing their differentiation from unmodified species during the oxidation step. The specific recovery after oxidation of miRNAs such as miR-7 and miR-142-3p from MIME-expressing Jurkat T cells further validates the sensitivity of this approach. The ability to distinguish between exogenous miRNAs (produced by human Jurkat T cells) and endogenous miRNAs (produced by mouse MIN6B1 cells), is further demonstrated by the differential recovery of Hsa-miR-155 and Mmu-miR-155, underscoring the specificity of the MIME-seq 2.0 method for tracking RNA transfer across cell types. Moreover, the lack of oxidation protection for U6, a non-Argonaute-interacting RNA, further highlights the specificity of the method for sncRNAs associated with Ago proteins.

The application of MIME-seq 2.0 to track miRNA transfer was further validated in C2C12 myotubes, where miR-1 and miR-133b, two myogenesis-related miRNAs [12], were detected in MIN6B1 cells following exposure to EVs. These results provide further validation of the ability of MIME-seq2.0 to track miRNA transfer between different cell types, suggesting its potential for monitoring miRNA transfer in vivo. This aspect of the study is particularly promising for understanding how miRNAs contribute to tissue-specific gene regulation, immune cell communication, or the spread of disease-associated miRNAs across different cell populations [5]. Despite the advantages presented above, the use of MIME-seq 2.0 presents some limitations. First, evidence that a sncRNA is closely associated with Ago proteins in living cells obtained by MIME protection, does not allow to conclude that it functions as a miRNA. Indeed, not all sncRNAs binding to Ago2 display miRNA-like repressive activities [3]. Second, although the MIME-seq 2.0 method can be adapted to efficiently monitor the transfer of sncRNAs between different cells, this technique cannot be applied to sncRNAs that are not methylated by the MIME enzyme, such as piRNAs and most tRF. Moreover, the sensitivity of the MIME-seq2.0 approach may not enable the detection of low-abundance miRNAs or the transfer of miRNAs that are already highly abundant in the receiving cells.

In summary, MIME-seq 2.0 represents a robust, innovative method for studying binding of small RNAs to Argonaute protein. This approach offers clear advantages over conventional techniques by providing a sensitive and specific means of identifying oxidized versus non-oxidized RNA species and by enabling the evaluation of RNA–protein interactions. Furthermore, the ability to track RNA transfer via EVs in different cell models expands the potential applications of this technique, including the study of intercellular communication and RNA-based therapies. Given its broad applicability, MIME-seq 2.0 holds great promise for advancing our understanding of small RNA biology and RNA-mediated cellular processes.

## STATEMENTS & DECLARATIONS

## Acknowledgement

We are very grateful to Drs. Luisa Cochella and Ariane Mandlbauer from the John Hopkins University of Baltimore for generously providing the plasmids for MIME expression and for the advice on the MIME-seq technique.

## DATA AVAILABILITY

The datasets generated during this study are available in the GEO repository under the accession number GSE313744 (token for reviewers upwtwyoybnutpuh).

